# High-Sensitivity Analysis of Native Bacterial Biofilms Using Dynamic Nuclear Polarization-Enhanced Solid-State NMR

**DOI:** 10.1101/2024.09.25.614951

**Authors:** Chang-Hyeock Byeon, Ted Kinney, Hakan Saricayir, Kasper Holst Hansen, Faith Scott, Sadhana Srinivasa, Meghan K. Wells, Frederic Mentink-Vigier, Wook Kim, Ümit Akbey

## Abstract

Bacterial biofilms cause persistent infections that are difficult to treat and contribute greatly to antimicrobial resistance. However, high-resolution structural information on native bacterial biofilms remain very limited. This limitation is primarily due to methodological constraints associated with analyzing complex native samples. Although solid-state NMR (ssNMR) is a promising method in this regard, its conventional applications typically suffer from sensitivity limitations, particularly for unlabeled native samples. Through the use of Dynamic Nuclear Polarization (DNP), we applied sensitivity enhanced ssNMR to characterize native *Pseudomonas fluorescens* colony biofilms. The increased ssNMR sensitivity by DNP enabled ultrafast structural characterization of the biofilm samples without isotope-labelling, and chemical or physical modification. We collected 1D ^13^C and ^15^N, and 2D ^1^H-^13^C, ^1^H-^15^N and ^13^C-^13^C ssNMR spectra within seconds/minutes or hours, respectively which enabled us to identify biofilm components as polysaccharides, proteins, and eDNA effectively. This study represents the first application of ultrasensitive DNP ssNMR to characterize a native bacterial biofilm and expands the technical scope of ssNMR towards obtaining insights into the composition and structure of a wide array of *in vitro* and *ex vivo* biofilm applications. Such versatility should greatly boost efforts to develop structure-guided approaches for combating infections caused by biofilm-forming microbes.

## Introduction

Bacteria form structured communities called biofilms, which create a protective environment for the encased cells against diverse stressors. Biofilm-associated bacteria are responsible for approximately 80% of chronic infections^[1]^ and significantly contribute to the rise of antimicrobial resistance (AMR).^[2]^ Biofilm cells display antibiotic tolerance levels that are several orders of magnitude higher than free-living planktonic cells.^[3]^ Diseases linked to AMR cause around one million deaths annually and could surpass cancer as the leading cause of death by 2050.^[4]^ Addressing the critical issues of biofilm mediated infections and antimicrobial resistance requires development of innovative therapeutic strategies that specifically target biofilms. These strategies should focus on inhibiting biofilm formation without harming human cells, employing interventions directed at specific biofilm structural components.^[5]^

The structural integrity of biofilms relies on a complex extracellular matrix (ECM), comprising polymeric proteins, polysaccharides, lipids, extracellular DNA (eDNA), and other metabolites.^[6]^ Disruption of key structural elements within biofilms, such as functional amyloids or polysaccharides, leads to biofilm destabilization, making the cells more susceptible to drug interventions.^[2a]^ Despite the critical role of these components, there remains a notable lack of structural information. For instance, there are only few structures currently available for the biofilm forming functional bacterial amyloids (FuBA): low-resolution structure of CsgA from *E. coli*, high-resolution structures of TasA from *B. subtilis*, and high-resolution structure of PSMα1 from *S. aureus*. ^[7]^ Similarly, high-resolution structural data on the biofilm polysaccharide fraction of ECM is greatly lacking.^[8]^ Addressing these knowledge gaps is vital for advancing our understanding of biofilm-related chronic infections and AMR. Structure-guided approaches are essential to drive significant progress in combating biofilm-related challenges, but they require high-resolution structural information.

Analyzing the structure of biological compounds within their native physiological environments presents significant technical challenges.^[9]^ NMR spectroscopy and cryo-electron microscopy/tomography are promising high-resolution techniques for investigating such complex biological systems.^[9-10]^ NMR spectroscopy offers the distinct advantage of providing a comprehensive, cumulative view of a biofilm sample at once. However, analyzing biomolecules within complex samples often requires the use of multidimensional (nD) NMR methods to resolve and simplify the observed signals, while isotopic enrichment at high concentrations is typically necessary for effective analysis. Magic-angle spinning (MAS) ssNMR (MAS NMR) can provide high-resolution structural insights on insoluble species, albeit with a sensitivity limitation. The lower sensitivity of ssNMR is problematic when isotopic enrichment is not feasible, such as for native biofilms extracted directly from patients with cystic fibrosis (CF).^[11]^ Enhancing sensitivity is thus imperative for rapid MAS NMR characterization of such important biological native samples. Sensitivity-enhanced dynamic nuclear polarization (DNP NMR) has been utilized to overcome sensitivity limitation, which has shown promising effectiveness in analyzing complex biological samples, including whole cells and extracts directly within their native environments.^[12]^

Over the past decade, MAS NMR has been employed to characterize the chemical composition of diverse biological samples, including bacterial, plant, and fungal cell walls. However, these studies have predominantly relied on 1D MAS NMR spectroscopy,^[12j, 13]^ which limits quantification due to inherent low resolution and signal overlap. Nonetheless, such 1D MAS NMR studies yielded valuable structural and compositional information on bacterial and fungal biofilms. ^[13-14]^ Multidimensional (nD) MAS NMR alleviate limitations associated with 1D MAS NMR spectroscopy, enabling the differentiation of signals originating from distinct components within complex biological mixtures. However, 2D NMR spectroscopy, in particular 2D ^13^C-^13^C correlations, is challenging compared to 1D ssNMR, and either requires isotope labelling or sensitivity-enhancement such as DNP. In this regard, a few examples of 2D MAS NMR applied to plant and fungal cell walls has been demonstrated with or without DNP.^[15]^ At the bacterial front, however to our current knowledge, only a few examples of nD MAS NMR experiments have been reported on natural abundance or isotope-labeled whole-bacteria, as well as extracted pure cell wall.^[16]^ Conventional high-resolution ^1^H- and ^13^C-detected MAS NMR have been employed on bacterial and plant cell walls in several investigations. ^[16b, 16c, 17]^ Recently, we and others showed the 2D ^1^H-^13^C MAS NMR by proton or carbon detection on nontuberculous mycobacteria (NTM) and *Pseudomonas fluorescens*.^[16a, 18]^ Finally, the high-sensitivity DNP NMR was utilized to characterize whole-cell or extracted cell walls from bacteria and yeast. ^[12j, 16c, 16d, 17, 19]^

Expanding the application of nD MAS NMR would greatly improve NMR-based structural characterization of native bacterial biofilms. Our primary objective is to address this information gap by employing hyperpolarized 1D/2D DNP NMR to extract detailed structural information about bacterial biofilms. In two recent studies, we utilized conventional room-temperature 1D and 2D MAS NMR spectroscopy to characterize colony biofilm samples of *Pseudomonas fluorescens* Pf0-1 and planktonic nontuberculous mycobacteria (NTM) at natural abundance.^[18, 20]^ We successfully identified over hundred distinct chemical sites in NTM planktonic whole-cells by utilizing high-resolution INEPT-based 2D MAS NMR and peak deconvolution. In this study, we demonstrate ultrasensitive 1D and 2D DNP NMR to characterize native colony biofilms of *P. fluorescens* Pf0-1. This biofilm model system has been previously demonstrated to produce diverse ECM components and employs a non-pathogenic bacterial strain advantageous to safe laboratory experimentation. Our findings showcase the promise of DNP NMR to acquire high-sensitivity 1D NMR spectra within seconds/minutes and 2D NMR spectra within hours from native, natural abundance bacterial biofilm samples. By combining ^13^C detection with 2D spectroscopy, we maximize spectral resolution, quantify different biofilm components, and gain new structural insights. The low temperature (∼100 K) required for DNP NMR allows simultaneous observation of both rigid and mobile ^13^C signals, owing to the favorable freezing of the sample. To the best of our knowledge, this study represents the first demonstration of sensitivity-enhanced DNP ssNMR to characterize an intact native bacterial biofilm. The experimental pipeline outlined here holds significant promise for advancing the technical boundaries of NMR-based biofilm research, as it can be readily applied to diverse biofilm preparations and model systems.

## Results and Discussion

### Structural characterization of native bacterial biofilm with hyperpolarized DNP ssNMR

Biofilm samples for MAS NMR analysis were prepared in two different ways, Figure 1. First, the native wet *P. fluorescens* colony biofilm was measured without any treatment, ‘wet biofilm’. Second, the biofilm was gently dried at 50°C, ‘dry biofilm’, effectively eliminating excess hydration. The drying step significantly improved sample packing efficiency by approximately tenfold, leading to increased signal-to-noise per unit time (SNT) with no effect on the CP-based spectra compared to the wet biofilm sample, Figure 2A,B, as we also showed recently by conventional ssNMR.^[18b]^ Both wet and dry biofilm samples were incubated with radical for DNP hyperpolarization and measured at ∼100 K with μw irradiation, Figure 1. We employed the AsymPol-POK radical as the DNP polarization agent dissolved in a DMSO/water (10:90% v:v) DNP buffer, which has demonstrated high efficiency at higher magnetic fields with short polarization buildup times at 10 mM concentration. The NMR and EPR characterization results are shown in Figure 2A-E. Moreover, the EM micrographs of the wet and dry biofilm samples were recorded prior to the incubation with radical, which indicate intact bacteria surrounded by ECM, Figure 2F.

**Figure 1.**
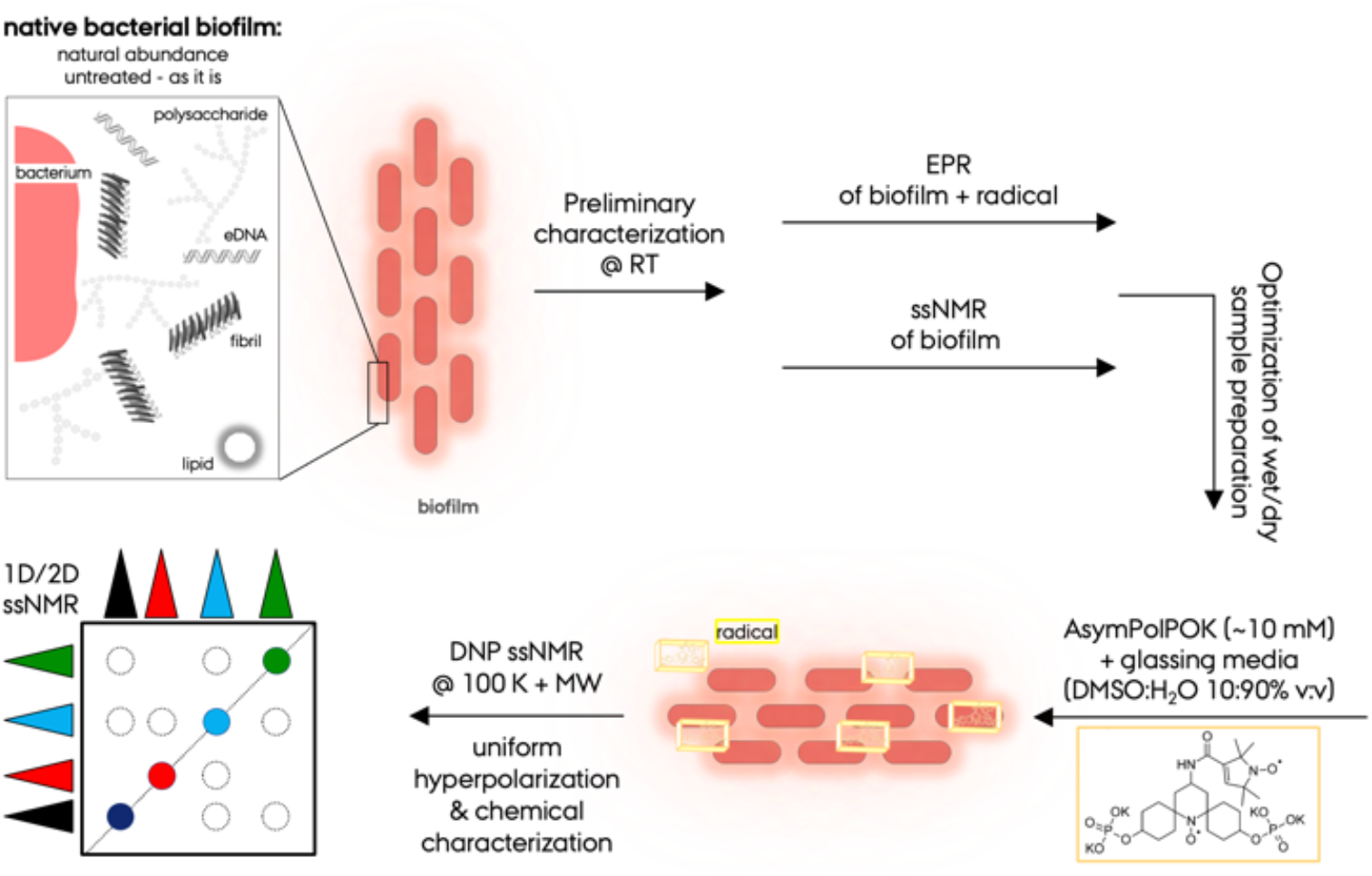
Experimental protocol for the hyperpolarized DNP NMR based structural characterization of native bacterial biofilms. The cartoon representation essential structural components of biofilm such as polysaccharides, fibrillar functional amyloid proteins, eDNA and lipids. From the native biofilm sample preparation towards the final ssNMR spectrum, Experimental conditions require careful optimization to maximize the sensitivity and reproducibility by including MAS NMR and EPR characterization.

**Figure 2.**
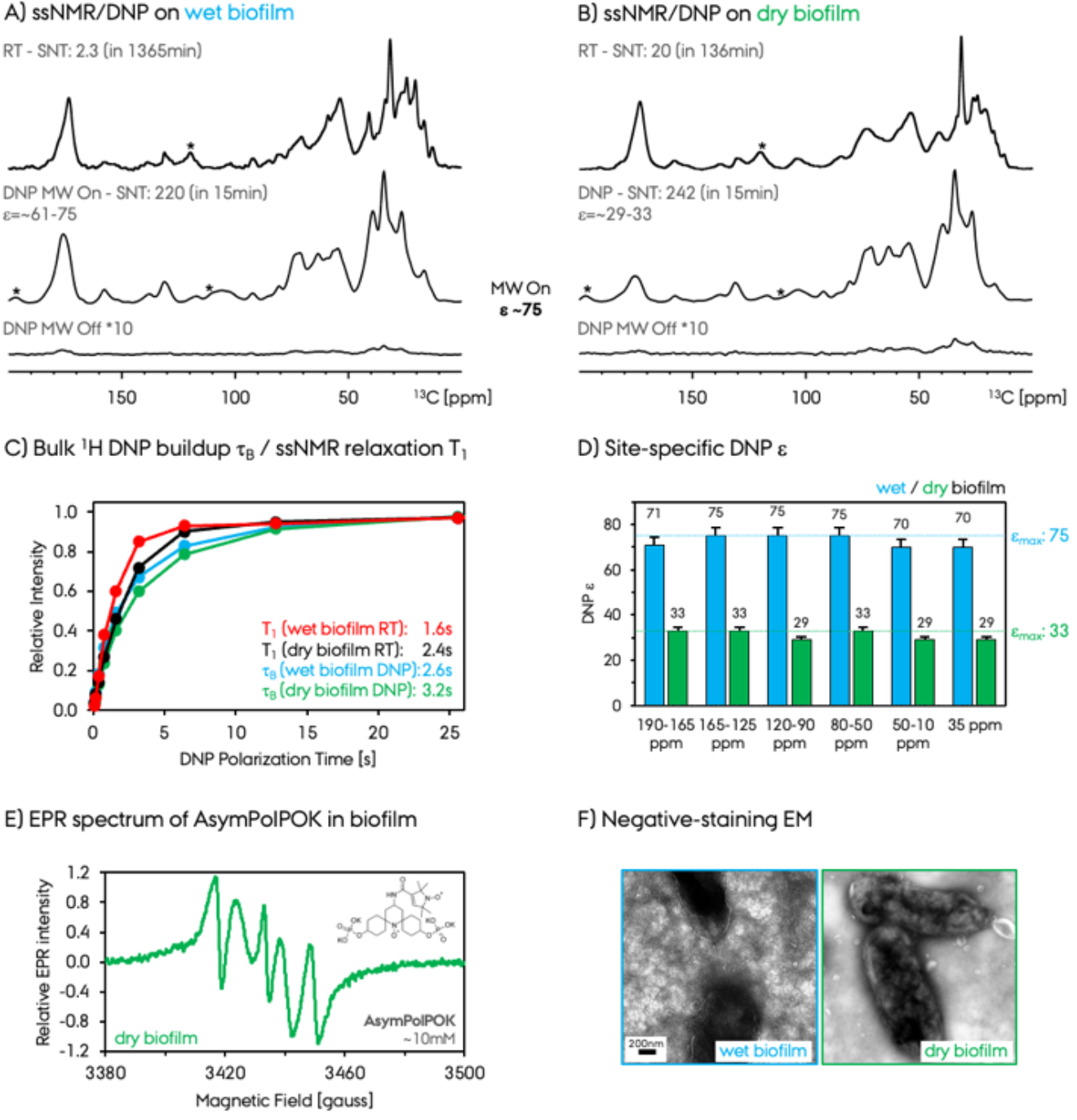
The 1D ^13^C CPMAS spectra were recorded with the conventional ssNMR at ∼300 K, 10 kHz and 750 MHz magnet, and the high-sensitivity DNP-enhanced ssNMR at ∼100 K, 10 kHz and at 600 MHz magnet on A) wet (native) and B) dry biofilm preparations. The signal to noise ratio per unit time (SNT, determined by (S/N)/(minute^0.5^)) values are given for a direct sensitivity comparison along with the total experiment times. For the DNP ssNMR spectra the μw on and off spectra are shown. The maximum DNP enhancements for the wet and dry biofilm samples are σ=75 and 33, respectively. C) The DNP buildup times (τB) and the ssNMR relaxation times (T_1_) are recorded by saturation-recovery method. The normalization was done at 52.8 s to 1.0 for each data set separately. D) Site-specific DNP σ for wet and dry biofilm samples. 190-165 ppm indicates CO signals, 165-125 ppm aromatic and nucleic acids, 120-90 ppm polysaccharides, 80-50 ppm polysaccharides and Cα, 50-10 ppm Cα and aliphatics, and ∼35 ppm lipids. Error bars represent a constant 5% error. E) The EPR spectrum of the freshly prepared dry biofilm sample containing 10 mM AsymPol-POK radical recorded at room temperature right before performing the DNP experiment. The ^13^C peaks at ∼119.7 ppm for room-temperature spectra and ∼111.4 ppm for DNP spectra are spinning sidebands and marked with asterisks. F) The negative-staining EM micrographs of the wet and dry biofilm samples. The scale bar is 200 nm.

DNP hyperpolarization resulted in a large signal enhancement of ε:∼75 for the wet biofilm preparation in the ^13^C CPMAS DNP-NMR spectra, Figure 2A and 3A. This represents one of the largest DNP enhancement observed among various native sample preparations on a 600 MHz DNP system, greatly facilitating an NMR based structure characterization. In previous studies, DNP enhancements of approximately ε:∼45-60 were observed for whole cell preparations (at 600 MHz), ε:∼25 for a drug inside intact cells (at 400 MHz), ε:∼30 for cell wall of whole or disrupted bacteria (at 400 MHz), and ε:∼30 for fungi/plant samples where the glycerol and/or polysaccharide fraction exhibited a larger enhancement of ε:∼90 for *Aspergillus fumigatus* fungus (at 600 MHz).^[12h, 19b, 21]^

In contrast to the large DNP enhancement observed for the wet biofilm, the dried biofilm preparation protocol with impregnation resulted in a smaller enhancement of ε:∼33 and a longer DNP buildup time, Figure 2B-D. These differences are attributed to a non-uniform biradical distribution and potentially increased microwave absorption by the sample. Moreover, variation in the DNP enhancements were observed for sample preparations with different radical incubation times, and we observed a decay in DNP enhancement at longer incubation times at room temperature (data not shown). This is due to a possible degradation of the radical within the native biofilm environment, as shown by EPR spectra showing deactivated AsymPol-POK after prolonged incubation, as seen in Figure SI1. This phenomenon was previously reported in other *ex vivo* DNP sample preparations in mammalian cells. ^[22]^ The data presented in this study were thus acquired from freshly prepared biofilm samples and measured immediately. These results highlight the importance of careful optimization of the sample preparation process to maximize the DNP effect in native biofilms.

The large signal improvements confirm the significant potential for characterizing native biofilms. The acquisition of 1D ^13^C MAS NMR spectra is accomplished in seconds/minutes using DNP NMR, thereby yielding a signal-to-noise ratio (SNT) comparable to that typically require hours/days by conventional MAS NMR, Figure 2A,B. The signal enhancement of ε∼75 results in a potential reduction in the time required for MAS NMR experiments by a factor of approximately 5625-fold (75^2^). Similarly, a signal enhancement of ε∼33 as observed in the dry biofilm sample, can reduce total experiment time by approximately 1000-fold (33^2^). The larger packing efficiency for dry biofilm sample increases its apparent SNT despite lower DNP enhancement. This sensitivity enables ssNMR experiments that were previously impractical due to extended acquisition times. A SNT: 220 is obtained at 1D ^13^C CPMAS DNP NMR spectrum for the wet biofilm sample, ∼100-fold larger compared to the conventional CPMAS spectrum recorded at room temperature per unit time, Figure 2A. For the dry biofilm sample, a SNT: 242 is obtained via DNP NMR, ∼12-fold larger compared to the CP MAS NMR spectrum due to the lower enhancement of this sample, Figure 2B. Despite the lower DNP enhancement observed for dry sample, the overall SNT is larger compared to the wet sample due to increased packing.

The bulk proton DNP buildup times (τB) were measured with μw irradiation at 100 K for both samples and compared to the bulk proton T1 relaxation times recorded at ∼300 K, Figure 2C. The τB values are only slightly longer than the room temperature T1 values, due to the favorable rapid buildup behavior of the AsymPol-POK radical.^[23]^ The electron paramagnetic resonance (EPR) spectrum of AsymPol-POK in freshly prepared native wet biofilm sample is shown, Figure 2E. These relatively short τB times significantly improve the overall SNT, especially in contrast to the buildup times observed when using different radicals as AMUPOL.^[24]^ The dry biofilm sample exhibited longer τB/T1 ratios compared to the wet sample. An attempt was made to quantify the differences in DNP τB times for various chemical sites within the wet and dry biofilm samples. Notably, the proton τB values determined for CO, nucleic acids, aromatics, polysaccharides, and Cα sites were found to be very similar (Figure SI2). This suggests a uniform distribution of the radical within the biofilm using the current sample preparation protocol, Figure 2D. Using the tentative ^13^C assignments in our previous MAS NMR study at room temperature, we estimated uniform DNP enhancements ε:70-75 and ε:29-33 for different chemical sites for the wet and dry biofilm sample, respectively, Figure SI3.^[18b]^ This indicates that radical is uniformly distributed in the complex biofilm matrix for the current colony biofilm system and sample preparation method, facilitating more quantitative characterization. Previous DNP NMR studies on complex environments highlight site-specific DNP enhancements for different chemical groups within the biofilm.^[12h, 19b, 21]^

To extend the chemical site information obtained from the biofilms, we recorded ^15^N CP DNP NMR spectrum on the native wet biofilm sample, Figure 3B. The nitrogen has a lower gyromagnetic ratio and sensitivity compared to carbon nuclei, making it particularly benefit from DNP hyperpolarization. Remarkably, for the natural-abundance wet biofilm a high-sensitivity 1D ^15^N CP DNP NMR spectrum was recorded in just ∼35 minutes with SNT:11. For a comparison, the ^15^N CPMAS spectrum recorded at room temperature from a fully packed biofilm sample is shown in Figure SI 4, recorded in 273 minutes with a SNT:∼0.24. The SNT of the DNP enhanced ^15^N spectrum is ∼46-fold larger compared to the conventional room temperature MAS NMR. We performed tentative resonance assignments in the 1D ^13^C/ ^15^N DNP NMR spectrum, Figure 3B and Table 1.^[25]^

**Table 1:**
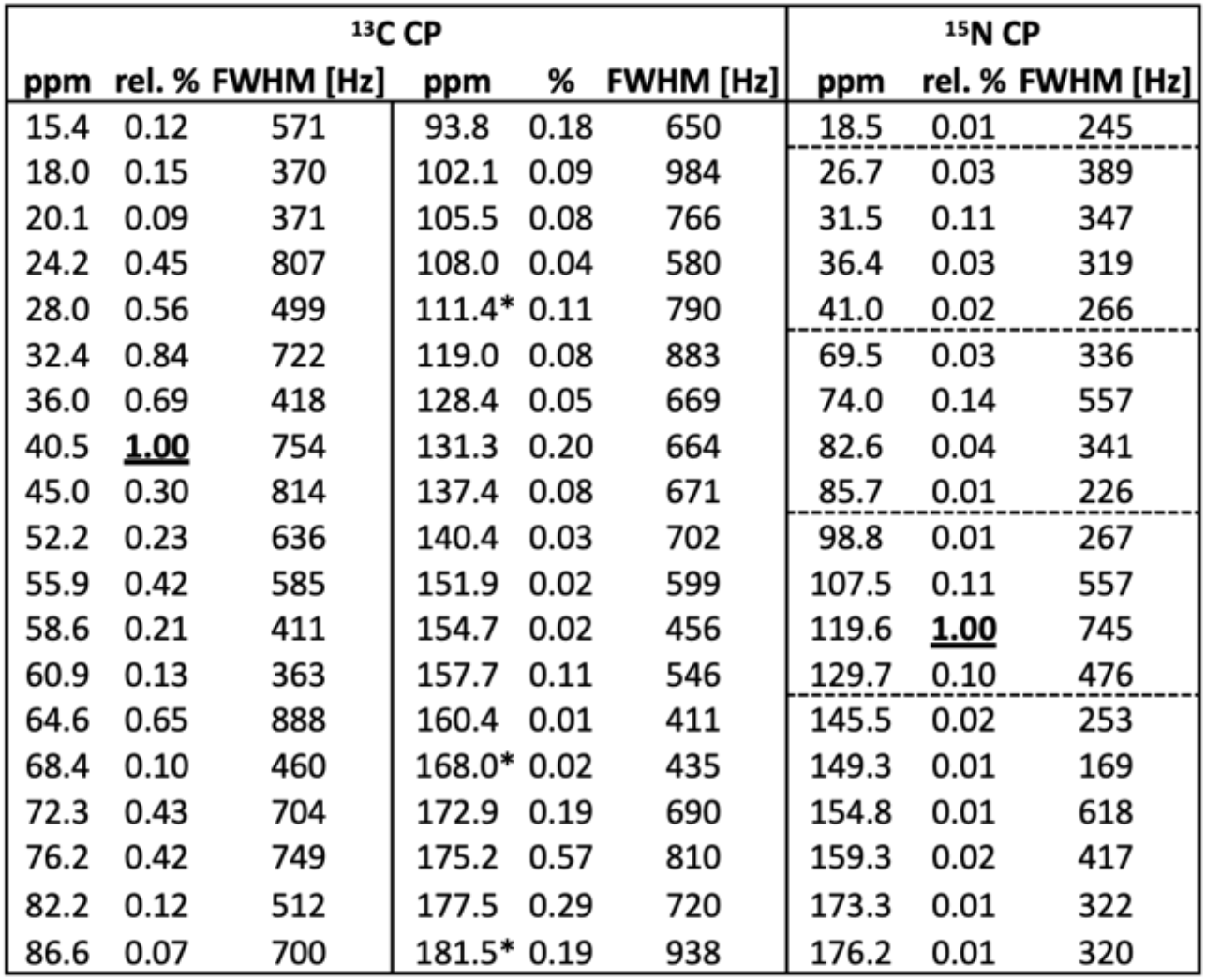
List of signals determined by deconvolution of the ^13^C and ^15^N CP DNP NMR spectra of native wet bacterial biofilm shown in Figure 2A,B. Chemical shifts, relative integral ratios normalized to the maximum integral (bold number as 1.00) within the ^13^C or ^15^N peak lists. The linewidths and the tentative resonance assignments are given for ^15^N resonances. The average FWHM for ^13^C and ^15^N signals are ∼640 (∼4.3 ppm) and ∼380 Hz (∼6.3 ppm), respectively, with the processing parameters given in the methods section. The spectra were recorded on a 600 MHz NMR spectrometer. The ^13^C peak at 111.4 ppm is a spinning sideband and marked with an asterisk.

**Figure 3.**
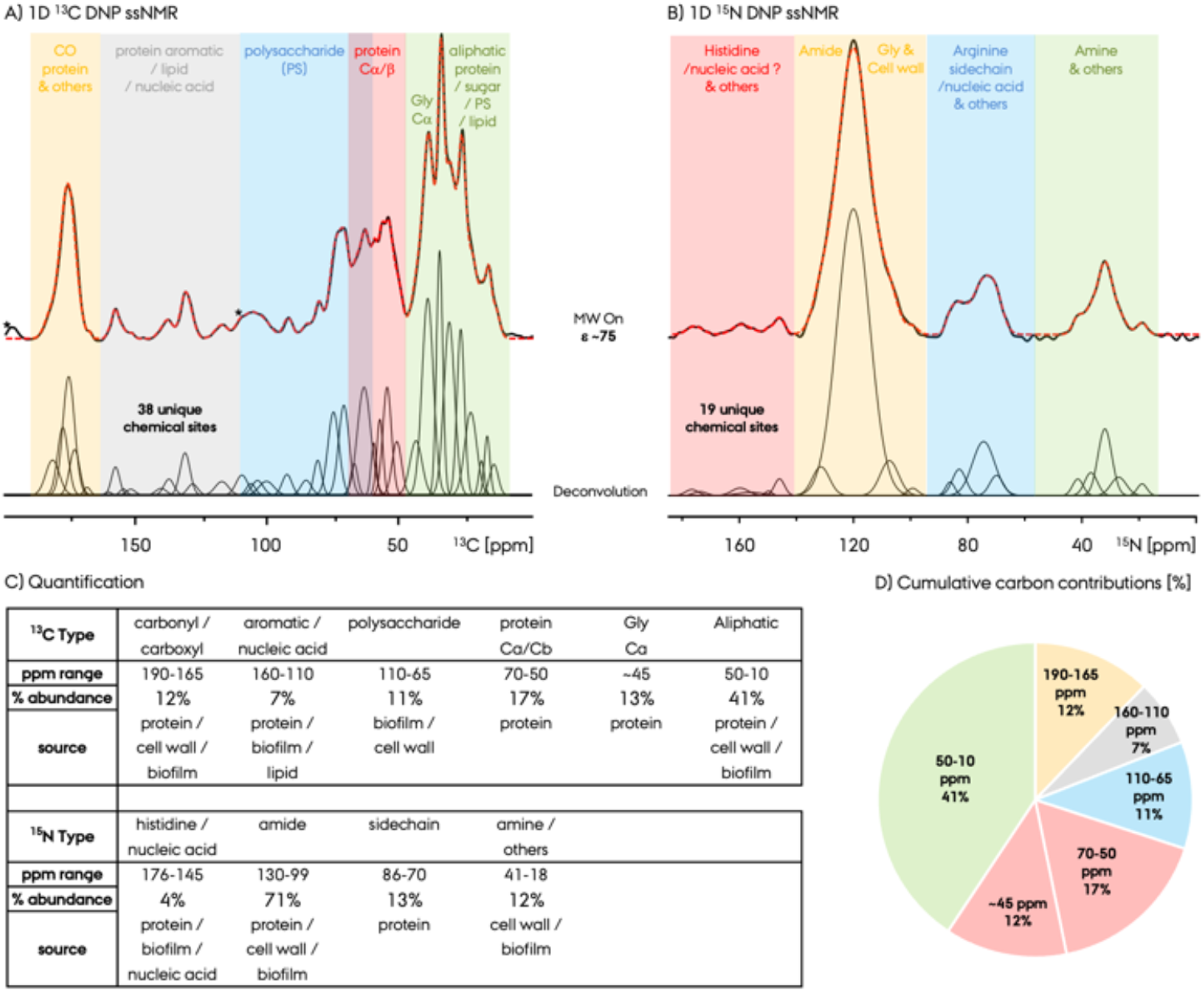
Quantification of different chemical sites in the A) ^13^C and B) ^15^N CPMAS DNP NMR spectra of natural abundance native wet bacterial biofilm. the long (1.2 ms for ^13^C and 0.4 ms for ^15^N) CP was used to ensure quantitative analysis. The individual peaks are determined by peak deconvolution utilizing ssNake program. The list of these peaks along with their linewidths are given in Table 1. Different tentative group assignments are color coded and labelled. The ^13^C peaks at ∼111.4 and ∼197 ppm are spinning sidebands and marked with asterisks in A, see Figure SI 5 for 1D ^13^C DNP ssNMR spectra recorded at different MAS.

The ^13^C full-width at half-maximum (FWHM) linewidths of ∼185 Hz (∼1 ppm) and ∼490 Hz (∼2.6 ppm) were observed at room temperature experiments for *Pseudomonas* biofilm sample based on INEPT and CP MAS NMR, respectively, on a 600 MHz NMR system. Notably, the linewidths obtained from the CP experiment were considerably broader compared to the mobile species observed in the INEPT spectrum. The room-temperature 1D ^13^C CP MAS NMR spectra for both wet and dry biofilm samples are shown in Figure 2A,B. Remarkably, lowering the experimental temperature from 300 K to 100 K for DNP NMR experiments yielded ^13^C CP spectra with similar resolution. The spectrum deconvolution indicates ∼30% resonance broadening at DNP NMR spectra at ∼100 K, and average linewidths of ∼640 Hz (∼4.3 ppm) and ∼380 Hz (∼6.3 ppm) were observed at ^13^C and ^15^N spectra recorded, respectively, Figure 3 and Table 1. These 1D DNP NMR based signal deconvolution combined with 2D NMR spectroscopy allows us to identify chemical sites in the *P. fluorescens* biofilm.

### High-throughput biofilm compositional analysis

The peak deconvolution of the 1D DNP NMR spectra identified 38 and 19 resonances for tentative assignments from the ^13^C and ^15^N CP spectra, respectively, Figures 3 and Table 1. The number of ^13^C peaks deconvoluted here are less compared to the 59 peaks obtained in the room temperature spectra, due to slight broadening of the resonances at ∼100 K. Correlated with the 2D ^1^H-^13^C and ^1^H-^15^N DNP NMR spectra shown in Figure 5, these peaks are consistent with the presence of proteins, polysaccharides and other species, and similarly observed in the previous studies on bacterial cells, cell wall, and biofilm.^[26]^

Following our recent room temperature biofilm study,^[18]^ we identified ^13^C signals in 1D/2D spectra from aliphatic carbons from proteins (Figure 2/3), carbohydrates, and lipids within the range of ∼10-50 ppm, as well as glycine Cα signals at around ∼45 ppm, peptide/protein Cα/Cβ signals at ∼50-70 ppm range, signals from polysaccharides between ∼65-110 ppm, aromatic signals from proteins and nucleic acids in the ∼110-165 ppm range, and carbonyl/carboxyl signals within the ∼165-190 ppm range. The presence of the NMR signal at around 175 ppm suggests the presence of proteins/peptides and polysaccharides, as this peak also disappeared in the room temperature INEPT spectrum due to the absence of protons at carbonyl/carboxyl carbons. NMR signals observed for alpha and aliphatic protein carbons within the range of approximately 10-70 ppm provides additional support. Moreover, in the 2D ^1^H-^13^C MAS NMR spectrum, these Cα carbon signals correlate with the appropriate alpha proton chemical shifts at ∼4.5 ppm, Figure 5.

A detailed deconvolution analysis of the carbonyl/carboxyl CO ^13^C signal region, between 165 and 190 ppm, was performed for the dry and wet biofilm samples recorded with both conventional MAS NMR and DNP NMR spectra, Figure 4. We quantified three distinct chemical sites at 172.9, 175.2 and 177.5 ppm. The peaks at 168.0 and 181.5 ppm are due to the spinning sidebands and despite their small abundance, helped to the accurate reproduction of the overall peak shape. By also taking advantage of the previous studies on bacterial cell walls, these resonances were tentatively assigned.^[26a, 26b]^ Accordingly, the 172.9 ppm signal was assigned to carbonyl site #1 from the protein, as well as to ester sites from the polysaccharides from the cell-wall/biofilm. 172.9 ppm was previously assigned to teichoic acid ester signals, but gram-negative *P. fluorescens* does not have teichoic acid or lipoteichoic acid. The 175.2 ppm signal was assigned to protein carbonyl site #2 predominantly from the biofilm protein fraction and the peptidoglycan peptides. Lastly, the 177.5 ppm signal was assigned to carboxyl signals from the uncross-linked ends of the cell wall peptidoglycans.

**Figure 4.**
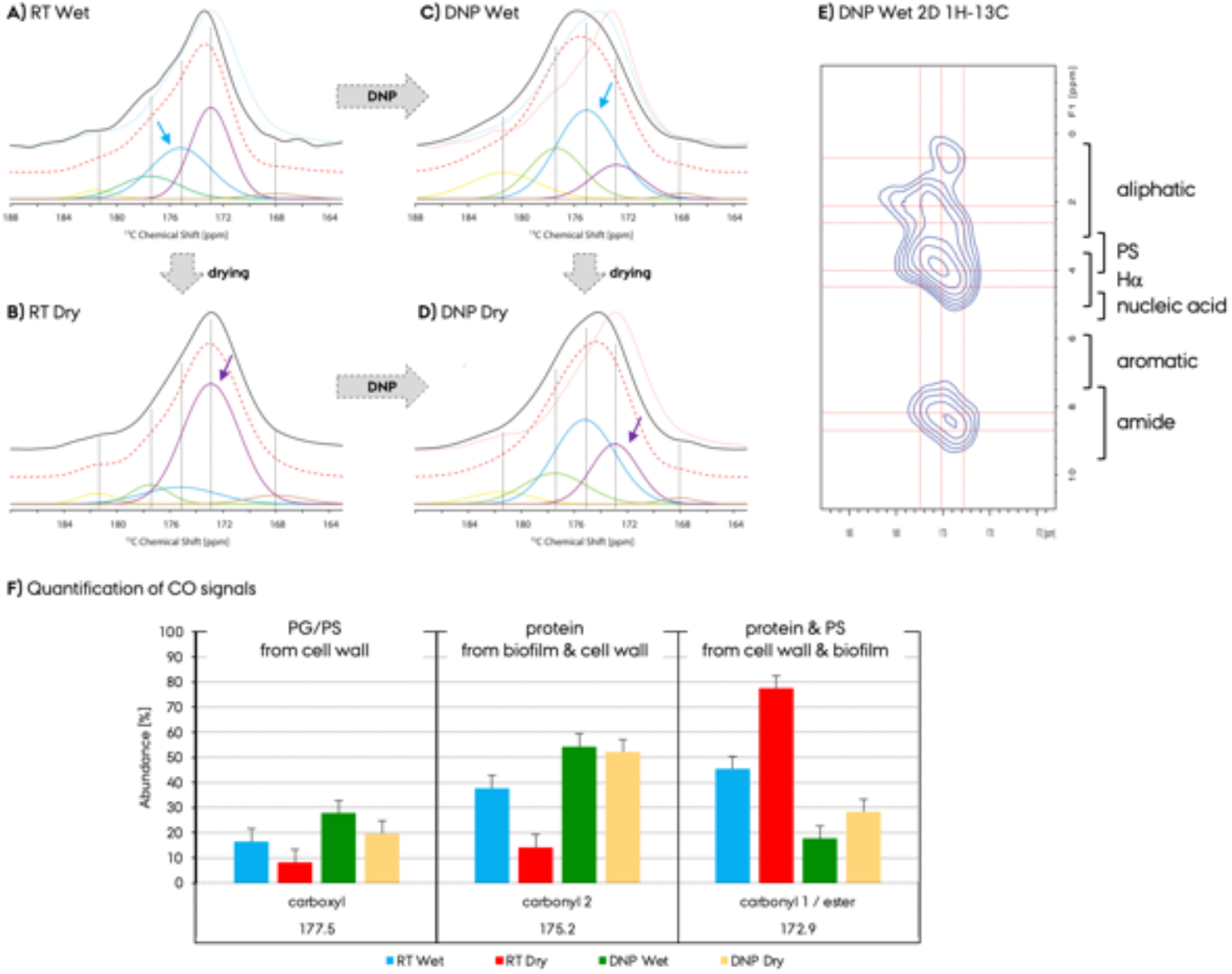
Quantification of the CO regions of the ^13^C CPMAS spectra recorded with A,B) conventional MAS NMR at room-temperature (RT) at ∼300 K and C,D) DNP NMR at ∼100 K. The CO signals for the ^13^C spectrum recorded for the A,C) wet and B,D) dry biofilm sample. E) The zoomed CO region of the 2D ^1^H-^13^C DNP NMR spectrum of the wet biofilm sample recorded with 100 μs CP contact time. F) The quantification of the three CO signals observed in the deconvolution analysis for wet/dry biofilm samples with constant 5% error bars. In A-D, the sum of the deconvoluted signals, the fitted spectrum, are shown as the spectrum with red dashed lines. Dashed black lines are placed at the corresponding chemical shifts of these five resonances. For direct comparison of the wet and dry spectra, the spectra from B and D are shown in A and C as light blue spectra. As a note, the color coding of the deconvoluted peaks and the bars in the graph are not correlated. The peaks at 168 and 181.5 ppm are due to spinning sidebands.

The 2D ^1^H-^13^C DNP NMR spectrum of the wet biofilm facilitates the group resonance assignment. Figure 4E, represents the proton correlations of these CO signals deconvoluted in Figure 4A-D. The proton dimension of the zoomed 2D DNP NMR spectrum in Figure 4E and 5 comprises chemical shifts specific to amides at ∼8-10 ppm, aromatics at ∼6-7 ppm, nucleic acids at ∼5 ppm, alpha at ∼4.5 ppm, sugar rings at ∼4 ppm, and aliphatics at ∼1-3 ppm. The 2D ^1^H-^13^C DNP NMR spectrum was recorded with short CP contact time (100 μs) and monitors directly bonded proton-carbon pairs. The 172.9 and 175.2 ppm peaks, e.g., correlate well with the amide and alpha protons at ∼8, ∼4.5 ppm, respectively, as well as with aliphatic protons, however, the 177.5 ppm peak has a much less amide proton cross peak and instead correlates with the carbohydrate proton signals at ∼4 ppm and aliphatic protons. The 175.2 ppm peak correlates with the aromatic protons at ∼6 ppm at lower contour levels (data not shown). This observation supports the tentative assignment by utilizing the previous bacterial whole cell and cell wall studies.^[16b, 27]^ The 172.9 and 175.2 ppm peaks are due to protein and/or polysaccharide species, whereas the 177.5 ppm peak is predominantly due to the cell wall peptidoglycan species.

**Figure 5.**
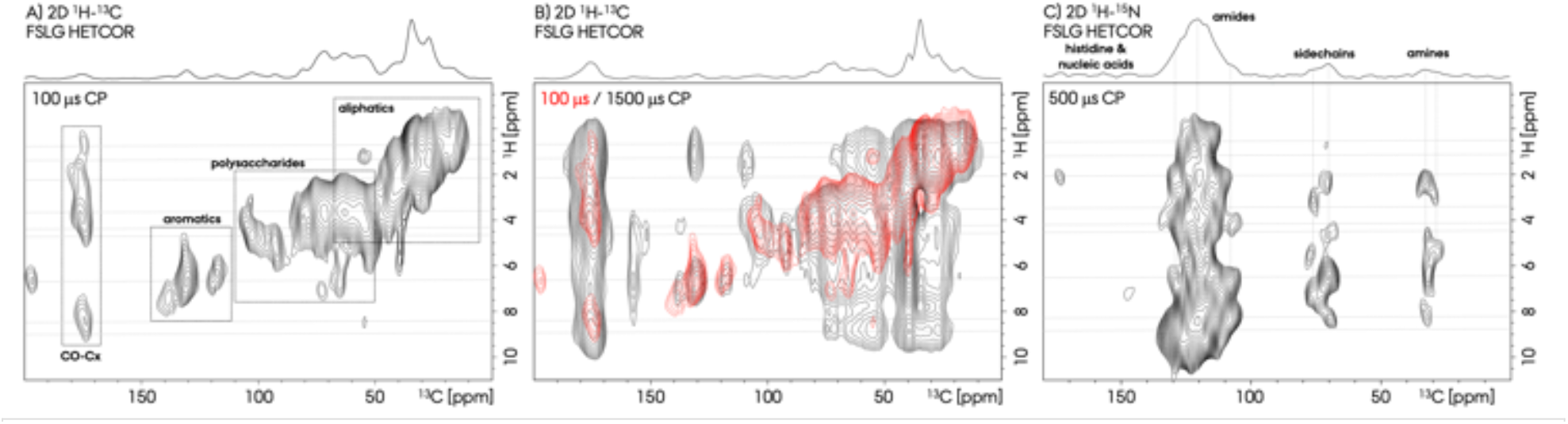
DNP-enhanced A,B) 2D ^1^H-^13^C and C) 2D ^1^H-^15^N FSLG HETCOR ssNMR spectra of natural abundance native wet bacterial biofilm. 100-1500 μs CP contact times were utilized. In B, the 2D spectrum with 100 μs contact time from A is shown in red, and the spectrum with 1500 μs is shown in black. The total experimental times are 18 minutes for A, 36 minutes for B, and 90 minutes for C.

The effect of drying the biofilm within the room-temperature (RT) and DNP ssNMR spectra were quantified by comparing the spectra in Figure 4A with B or Figure 4C with D. The relative peak integral percentage changes indicate that drying the biofilm increases the apparent abundance of the 172.9 ppm site relative to the others. This peak’s overall NMR ratio increased from ∼45% up to ∼78% up on drying (18% up to 28% for the DNP spectra), reducing the relative abundance of the 175.2 and 177.5 ppm sites to approximately half. This pattern indicates an increase in protein vs polysaccharide signal observation content due to the increased packing of biofilm sample. The DNP NMR spectra resulted in decreased relative abundance of the 172.9 ppm resonance for both dry and wet samples, but increased ratio for the other two resonances. In particular, the 175.2 ppm peak ratio in the CO region increased 1.4 and 3.7-fold for wet and dry biofilm samples, respectively. Overall, these observations indicate that DNP hyperpolarization enhances the signals from the protein/peptide fraction of the biofilm more relative to other fraction as PG.

The ^15^N CPMAS DNP ssNMR spectrum of the native wet biofilm composition was also analyzed for structural insights, Figure 3B. The ^15^N spectrum was recorded in 273 minutes (SNT: 0.24) by conventional NMR, whereas in 40 minutes (SNT:12) by DNP NMR, Figure SI4. The difference in the SNT is ∼50-fold between the conventional and DNP NMR, which approximates the DNP signal enhancement. The amide backbone nitrogen signals were observed at ∼120 ppm as the most abundant signal in the ^15^N spectra, originating from both protein, polysaccharide or peptidoglycan nitrogen sites within the biofilm. The relative abundance of the amide resonances is 71% of all the nitrogen signal integrated area and was deconvoluted into three significant resonances centered at around 107.5, 119.6 and 129.7 ppm, Figure 3B and Table 1. The signals at 107.5 and 129.7 ppm comprise ∼8% and 9% of the amide nitrogens, respectively, whereas the most prominent amide peak at 119.6 ppm comprises ∼82% of the total amides. The presence of these three amide resonances is similarly observed in the 2D ^1^H-^15^N DNP NMR spectrum, see dashed lines in Figure 5C. The resonance observed at 129.7 ppm compared to the major peak at 119.6 ppm indicates the amides at a different chemical environment due to different maxima, e.g., from biofilm versus from bacterial cell wall species.

In previous bacterial cell MAS/DNP NMR studies, two ^15^N amide resonances were observed which were assigned to less abundant peptidoglycan glycines at 107.5 ppm and to more abundant non-glycine amides from protein components at ∼120 ppm.^[28]^ In whole bacterial cell and cell wall studies, this glycine amide at 107.5 ppm was found to comprise ∼17-28% of all amides. We observe a much smaller abundance of ∼8% for this signal in the *P. fluorescens* biofilm sample. The lower relative abundance of the cell-wall/peptidoglycan glycine resonance compared to results from bacteria is consistent with the presence of large extracellular matrix surrounding the bacterial cells in the biofilm. In addition to amide resonances, amine (at 26.7-41 ppm) and protein sidechain (69.5-85.7 ppm) peaks also exist with relative abundance of 11% and 13% of all nitrogen signals, respectively. Moreover, the presence of resonances between 145-176 ppm (predominantly at ∼145.5, ∼159.3 and ∼176.2 with other minor ones listed in Table 1) indicates histidine and/or nucleic acid nitrogen signals,^[28a]^ which comprises ∼3% of the total nitrogen signal integrals. However, we did not observe any proton correlation from these cross-peaks in the 2D ^1^H-^15^N DNP NMR spectrum, most probably due to lower abundance and SNT.

### DNP facilitates resonance assignment by 2D ^13^C-^13^C correlation spectra

The 2D NMR spectrum is essential for quantitative spectral analysis, increases spectral resolution and aids resonance identification, particularly in regions with spectral overlap arising from proteins and polysaccharides in the biofilm. One of the most significant advancements enabled by the DNP sensitivity increase is the ability to record 2D DNP NMR spectra at natural abundance. Utilizing DNP enhancement of ε :∼75 we successfully recorded CP-based 2D ^1^H-^13^C (in 18/36 minutes, Figure 5A,B), 2D ^1^H-^15^N (in 90 minutes, Figure 5C), and 2D ^13^C-^13^C (∼17 hours, Figure 6) DNP NMR spectra. The conventional room temperature 2D ^13^C-^13^C MAS NMR spectrum of the dried sample recorded in ∼40 hours did not result in any cross peaks, Figure SI 6. The most demanding two-dimensional experiments is the 2D ^13^C-^13^C single-quantum – double-quantum (SQ-DQ) correlation experiment, due to the low natural abundance of ^13^C (approximately 1% of all carbons) and the extreme scarcity of ^13^C-^13^C pairs suitable for 2D detection (0.01% of all carbons). Nevertheless, we could record 2D spectrum using ∼50 mg of dry unlabeled biofilm sample. Drying the biofilm increased sample packing efficiency thus making the 2D ^13^C-^13^C spectrum possible to record within less than a day. Obtaining this spectrum from a wet biofilm was difficult due to the significantly less efficient rotor packing.

**Figure 6.**
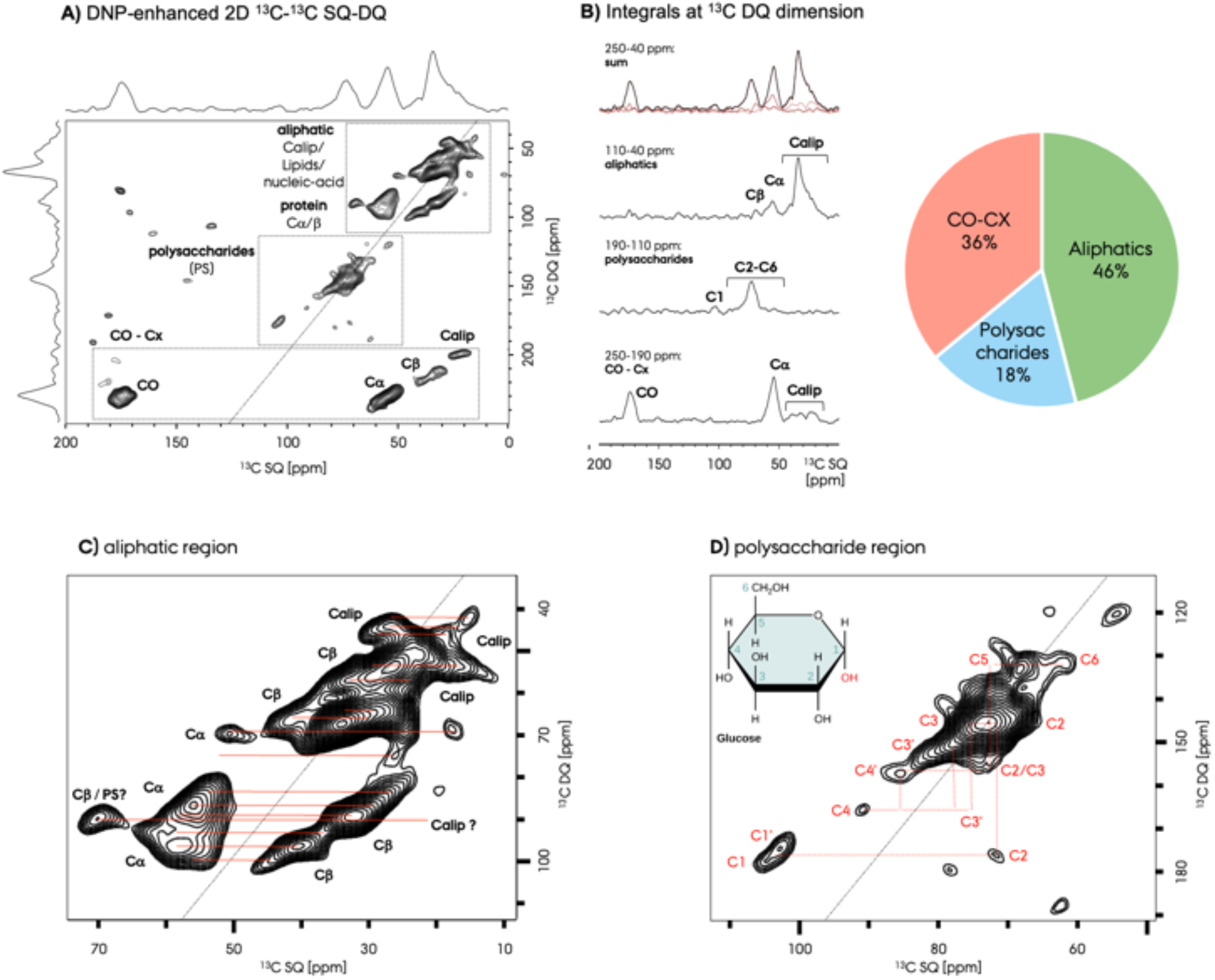
A) DNP-enhanced 2D ^13^C-^13^C refocused dipolar INADEQUATE SQ-DQ ssNMR spectrum of natural abundance native wet bacterial biofilm. 100-1500 μs CP contact times were utilized. B) 1D slices extracted between specific DQ chemical shift intervals, corresponding to the CO-Cx, polysaccharide and aliphatic (from protein, lipid, nucleic acid, and so on) chemical shift values. These 1D spectra highlights the potential of 2D ssNMR spectroscopy and results in the non-overlapped signals from those specific sites in 1000min./16.6h (3.25s x 288scans x 64). The zoom out regions for the C) aliphatic and D) polysaccharide chemical shift areas shown in A.

Tentative assignments obtained by utilizing 1D spectra are elaborated here by the cross-peaks observed in the 2D spectra with the additional proton chemical shift dimension for unambiguity in Figure 5. Different proton chemical sites are marked with dashed lines. We detected signals that can be attributed to the diverse components within the biofilm such as protons from amides, aromatics, nucleic acids, sugar rings, and aliphatics. For example, since the proton chemical shifts are different for protein and polysaccharide sites, the overlapping carbon chemical shifts in the 1D spectra can be identified more precisely as following the correlations of the carbon resonances with the proton dimension of the 2D ^1^H-^13^C MAS NMR spectra. These signals illuminate the interactions between different biofilm components including polysaccharides, lipids, and the cell wall. Particularly, the comparison of the short and long CP contact time 2D ^1^H-^13^C spectra gives hints for spatial proximities between different biofilm components, Figure 5A,B. The aromatic signals observed at ∼130 ppm ^13^C chemical shift correlates with a ∼6.5 ppm proton site at 100 μs CP contact time, Figure 5A. At longer CP contact time of at 1500 μs, there is an additional correlation from the protein aliphatic proton site but no polysaccharide proton correlation, Figure 5B indicating that the aromatic sites are not in close contact with biofilm polysaccharides. Additional indication of spatial segregation maybe the absence of amide proton correlation to any anomeric polysaccharide peak.

There are indications for the presence of nucleic acid signals in the 1D ^13^C and 2D ^1^H-^13^C DNP NMR spectra. In the 1D ^13^C spectra, the resonance observed at ∼93.8 ppm is specific to a nucleic acid carbon resonance, which correlates with a proton chemical shift at ∼5 ppm at short/long CP contact times in the 2D ^1^H-^13^C spectra. This proton resonance does not correlate with any of the polysaccharide, aromatic or protein Cα chemical shift supporting this tentative assignment. Nitrogen resonances observed above 140 ppm also indicates the presence of nucleic acids in the biofilm. The carbon chemical shift at 157.7 ppm could also be from nucleic acid signals since it correlates with the same proton signal at 5, and additionally with the 6.7 ppm proton peak at the 2D spectrum recorded with longer CP contact time.

The 2D ^13^C-^13^C SQ-DQ spectrum, Figure 6A, helps to identify resonances within the overlapped regions in the 1D ^13^C spectrum shown in Figure 3A. The overlapped regions can be resolved and analyzed in combination with the help from ^1^H-^13^C 2D spectra shown in Figure 5. Three distinct chemical shift areas are highlighted in Figure 6A, protein (aliphatic and Cα)/lipid/nucleic-acid, polysaccharide and carbonyl/carboxyl. As a result, chemical shift identification and specific integration of NMR resonances in the second dimension of a 2D ^13^C-^13^C SQ-DQ spectrum allows for the quantitative determination of different carbon pools corresponding to various chemical species, Figure 6B. The aliphatic and carbohydrate signal regions are zoomed out in Figure 6C,D. The cross peak locations and corresponding tentative resonance assignments are given that are obtained by utilizing the CCMRD carbohydrate and BMRB protein databases, as shown for other non-biofilm systems.^[12j]^

## Conclusion

We employed high-sensitivity DNP-enhanced ssNMR to characterize biofilm structure and composition *P. fluorescens* colony biofilm, with a particular focus on its protein and polysaccharide fractions. For the first time, we showcased the 2D MAS NMR spectra of a complex, native bacterial biofilm system without the need for purification and fractionation of individual biomolecules. These 2D DNP ssNMR spectra enable rapid observation of signals from the structural components of intact biofilms, which would otherwise not be possible. The findings from this study not only demonstrate the technical feasibility of DNP ssNMR to analyze diverse biofilm components, but they also set a solid foundation for future biofilm research. We acknowledge that the quantification of biofilm components via DNP may present challenges, necessitating further sample preparation refinements for each studied system. Furthermore, the availability of isotope-labeling provides an alternative avenue for 2D/3D DNP ssNMR experimental approaches, albeit not for patient derived samples. By introducing high-sensitivity and high-throughput structural DNP ssNMR characterization, our results aims to advance the field of structural analysis of biofilms and contribute to a deeper understanding of these complex biological systems.

## Acknowledgements

UA acknowledges financial support from University of Pittsburgh startup funding and the high-field NMR infrastructure at the Structural Biology Department, School of Medicine, University of Pittsburgh. WK was supported by funding from the National Institute of General Medical Sciences of the NIH 1R15GM132856. The National High Magnetic Field laboratory (NHMFL) is funded by the National Science Foundation Division of Materials Research (DMR-1644779 and DMR-2128556) and the State of Florida. A portion of this work was supported by the NIH P41 GM122698 and RM1-GM148766. F.J.S. acknowledges support from a postdoctoral scholar award from the Provost’s Office at Florida State University.

## Supplementary Information

### Experimental

#### Preparation of biofilm samples for NMR

The preparation of *Pseudomonas fluorescens* Pf0-1 colony biofilm samples was carried out as previously described._[20a, 20c, 29]_ Briefly, 20 μl of overnight liquid cultures were spotted and adsorbed onto solid *Pseudomonas* agar F (PAF) plates and incubated at ambient room temperature for three days. The resulting colony biofilms were then carefully scraped from the agar surface and stored in microfuge tubes at 4°C until MAS NMR analysis. For 13C-detected MAS NMR experiments, we employed thin-walled 3.2 mm rotors, with an approximate total biofilm material quantity of 40-60 mg. The transfer of the biofilm material into the rotor was executed utilizing a benchtop centrifuge, employing straightforward pipets directly affixed to the rotor, followed by centrifugation. It’s important to note that a considerable proportion of the native hydrated biofilm material consists of water, leaving a relatively small effective solid material content in the rotor. This was verified through a basic wet-biofilm drying. For the dry biofilm sample preparation, the wet biofilm was gently dried at 50°C in an oven under atmospheric conditions, effectively eliminating excess hydration and leaving a brittle and solid material. The drying process significantly increased the total material in the NMR rotor (∼10-fold), a consequence of the removal of excess hydration. Both wet and dried biofilm samples were analyzed in fully packed rotors.

#### DNP NMR Sample preparation and experiments

We tried two different sample preparations in terms of the biofilm treatment and radical incorporation. First, ∼50 mg of untreated native wet biofilm material with an effective solid material of ∼10-20% of total mass, was incubated with ∼20-30 μl of AsymPol-POK radical from a 10 mM stock radical solution in d6-DMSO:H_2_O (10:90%-v:v).^[23a, 30]^ Modest solvent deuteration was used since AsymPol-POK was shown to be efficient under such conditions._[30]_ The radical was added onto the wet biofilm slurry sample in three fractions. The resulting radical-absorbed biofilm mass was benchtop centrifuged into the NMR rotor. The second preparation protocol utilized a drying step for the biofilm material to remove the excess water, thus packing more solid material into the NMR rotor. For this purpose, the biofilm sample was dried at ∼50°C to yield a brittle solid. This powder was impregnated with ∼20-30 μL of AsymPol-POK, as in the first method, resulting in ∼30 mg of dry material packed into the NMR rotor. The presence of was verified by EPR measurements before each experiment to ensure the presence of sufficient active radical in the samples prior to measurements. The DNP enhancements were determined by direct comparison of μw on/off spectra. These sample preparation protocols demonstrate proof-of-concept principle but would benefit from further optimization to maximize DNP NMR sensitivity.

#### Conventional and DNP ssNMR Spectroscopy

All conventional room-temperature MAS NMR experiments were performed on a 750 MHz Bruker Avance III spectrometer, equipped with a low-temperature triple-resonance 3.2 mm probe. 10 kHz MAS spinning rate was applied with a set temperature of 275 K for all measurements, corresponding to ambient sample temperature. Pulses of 3.3 μs and 5 μs were employed for _1_H and _13_C nuclei, respectively. 1 ms contact time was utilized in the CP experiments with a 70-100% ramp on the proton channel. Proton dipolar decoupling of ∼90 kHz was applied throughout the acquisition. _1_H chemical shifts were directly referenced to 0 ppm, utilizing DSS as an internal standard incorporated into the NMR samples. The _13_C and _15_N chemical shifts were indirectly referenced by using the _1_H frequency._[31]_ The 2D _13_C-_13_C PDSD spectrum of the *P. fluorescens* sample from a fully packed 3.2 mm rotor was recorded in ∼200 hours with 20 ms mixing time. The spectrum was recorded at 275 K set temperature and 10 kHz MAS. 1024 scans were recorded with 200 indirect dimension increments and 1 second recycle delay.

The DNP NMR spectra were acquired on a 600 MHz Bruker NEO NMR spectrometer utilizing a 3.2 mm LT DNP probe at ∼100 K and 10 kHz MAS and using a sapphire rotor containing ∼30 mg of material. All the data were recorded at ∼100 K and 10 kHz MAS. The μw power was carefully optimized for each sample at the home-build quasi-optical mw setup at the NHMFL to achieve maximum signal._[32]_ Conventional room temperature 1D _13_C CP spectra were recorded with 43k (23 hours) and 8k (2.3 hours) scans and with 2 and 1 seconds recycle delays for the wet and dry biofilm samples, respectively. The 1D _13_C CP DNP NMR spectra were recorded with 256 scans with ∼3.5 seconds recycle delay for wet and dry samples, respectively, with/without μw irradiation in ∼15 minutes. 1D _15_N CP DNP NMR spectrum was recorded with 648 scans and 3.5 seconds recycle delay for wet biofilm in ∼35 minutes.

Signal to noise ratios, S/N, were determined within a signal region of 90 to 10 ppm and a noise region of -70 to -150 ppm for all spectra. The calculated S/N values were used to compute the signal-to-noise per unit time (SNT) obtained by dividing S/N by the square root square of the polarization time in minutes. For the _15_N SNT, calculation the amide signal is used. Spectral fitting and peak deconvolution were performed using the ssNake program package._[33]_ 1D spectra was processed by using gaussian function with 35 Hz broadening (LB: -35 and GB=0.02 in Topspin).

The 2D _1_H-_13_C FSLG CP MAS NMR spectrum was recorded with 1k scans for dry biofilm sample in ∼22 hours. 1 second recycle delay was used and a total of 78 indirect dimension data points were recorded with an increment of 100 μs. The 2D DNP NMR spectra were recorded at. The 2D _1_H-_13_C FSLG CP DNP NMR spectra were recorded with 100 μs CP contact times (probing primarily one-bond correlations) and 8 scans, or 1500 μs CP contact times (probing one-bond & medium/long correlations) and 16 scans, respectively, utilizing 30 indirect dimension increments. Total experiment time was 18 or 36 minutes. The 2D _1_H-_15_N FSLG CP DNP NMR spectrum was recorded with 500 μs CP contact time and 32 scans utilizing 40 indirect dimension increments._[34]_ A _1_H transverse RF field of 100 kHz was applied for the FSLG corresponding to an effective field strength of 122 kHz.

Total experiment time was 90 minutes. The 2D _13_C-_13_C SC-DQ SP-C5 DNP NMR spectrum was recorded with 500 μs CP contact time and 288 scans utilizing 64 indirect dimension increments._[35]_ Total experiment time was ∼17 hours (∼1000 minutes). The 2D spectrum was processed with gaussian broadening (with 35 or 50 Hz at GB = 0.02 in Topspin) for direct dimension and by a mixed sine squared for indirect dimension (SSB = 3 in Topspin).

#### EPR spectroscopy

The EPR spectra we collected on an EMX Nano at room temperature, using 0.1 mT of field modulation. The μw frequency was ∼ 9.4 GHz. The samples were packed into the 3.2 mm sapphire rotors and measured as is in a 4 mm EPR tube.

#### Negative-staining Electron Microscopy (EM)

400 mesh carbon film copper grids were glow discharged for 90 seconds. Dry and wet biofilm samples dispersed in ∼25 μL of 20 mM sodium phosphate solution at pH 7.8 and then applied to grids and left for 10 seconds before side blotting on filter paper. Grids were negative stained with 2% (weight/volume) uranyl acetate for 10 seconds before side blotting on filter paper. EM micrographs were recorded at a FEI Tecnai TF20 microscope with a field emission gun operating at 200 kV equipped with a TVIPS XF416 CMOS camera. No further image processing was done.

**Figure SI 1:**
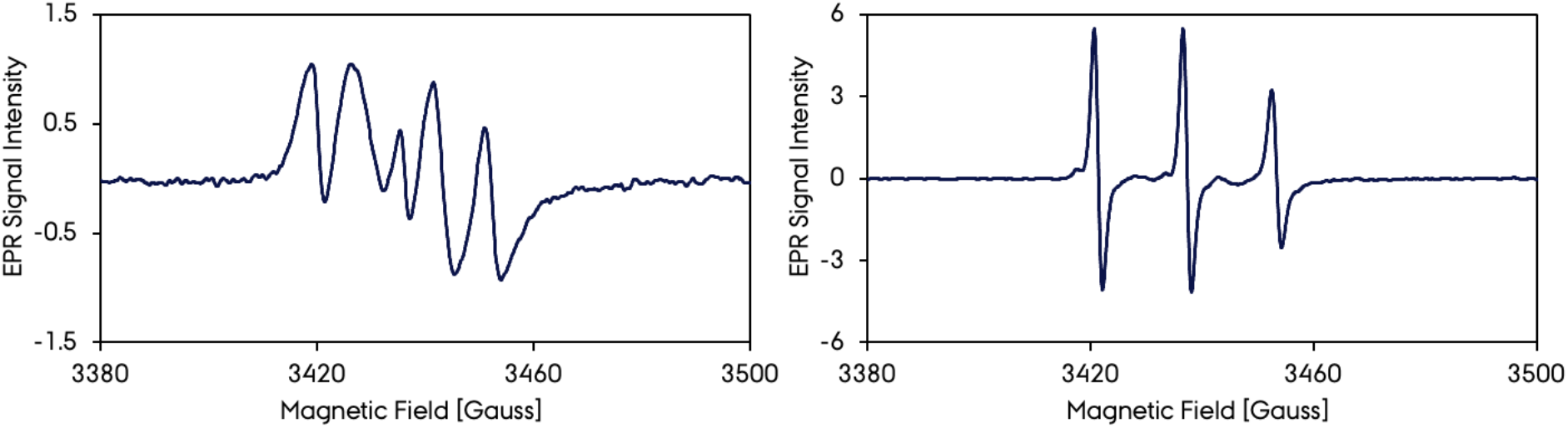
EPR of A) active and B) deactivate radical and biofilm sample.

**Figure SI 2:**
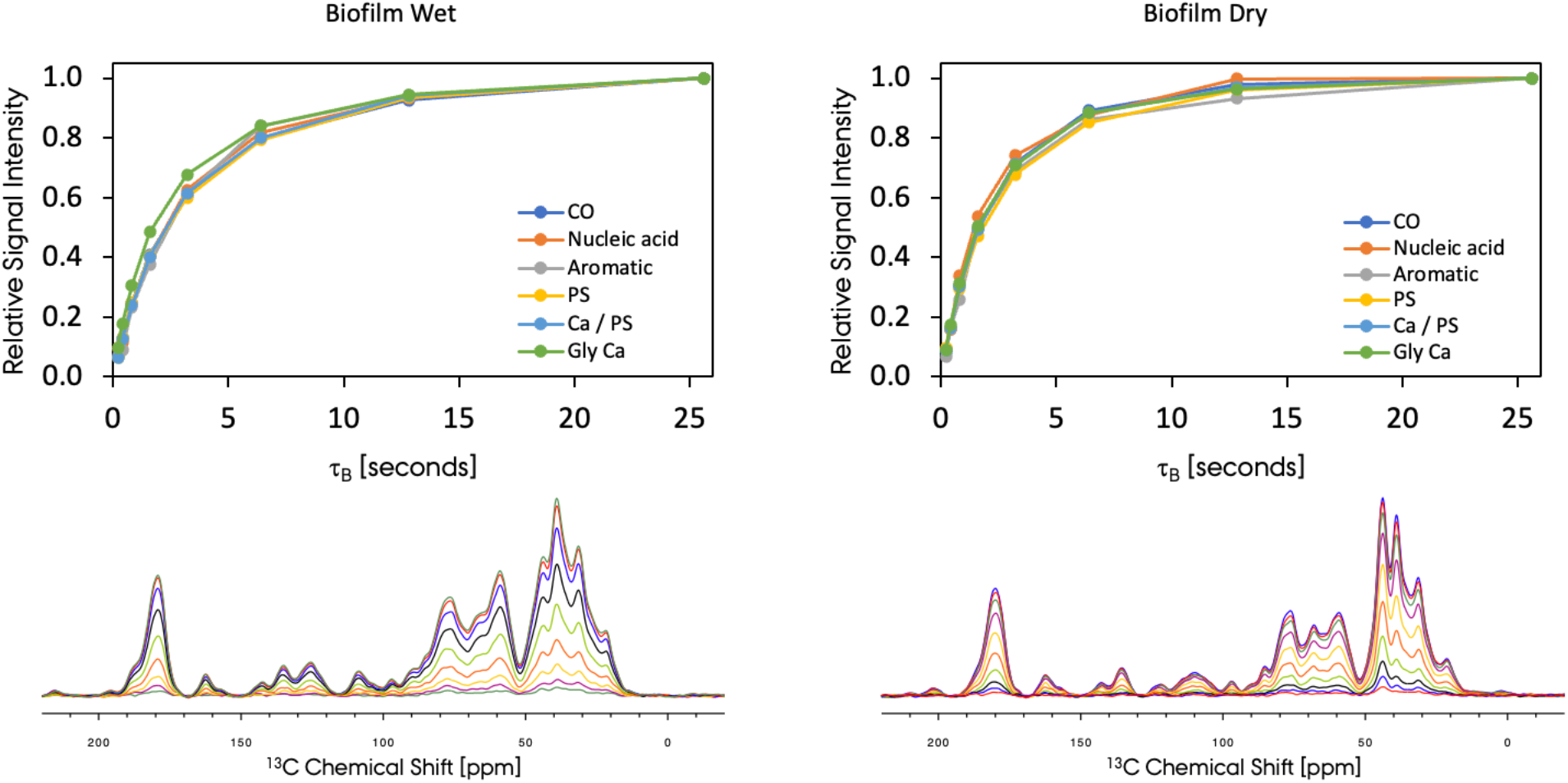
Site-specific τ_B_ times for wet and dry P. fluorescens biofilm samples recorded with DNP ssNMR.

**Figure SI 3:**
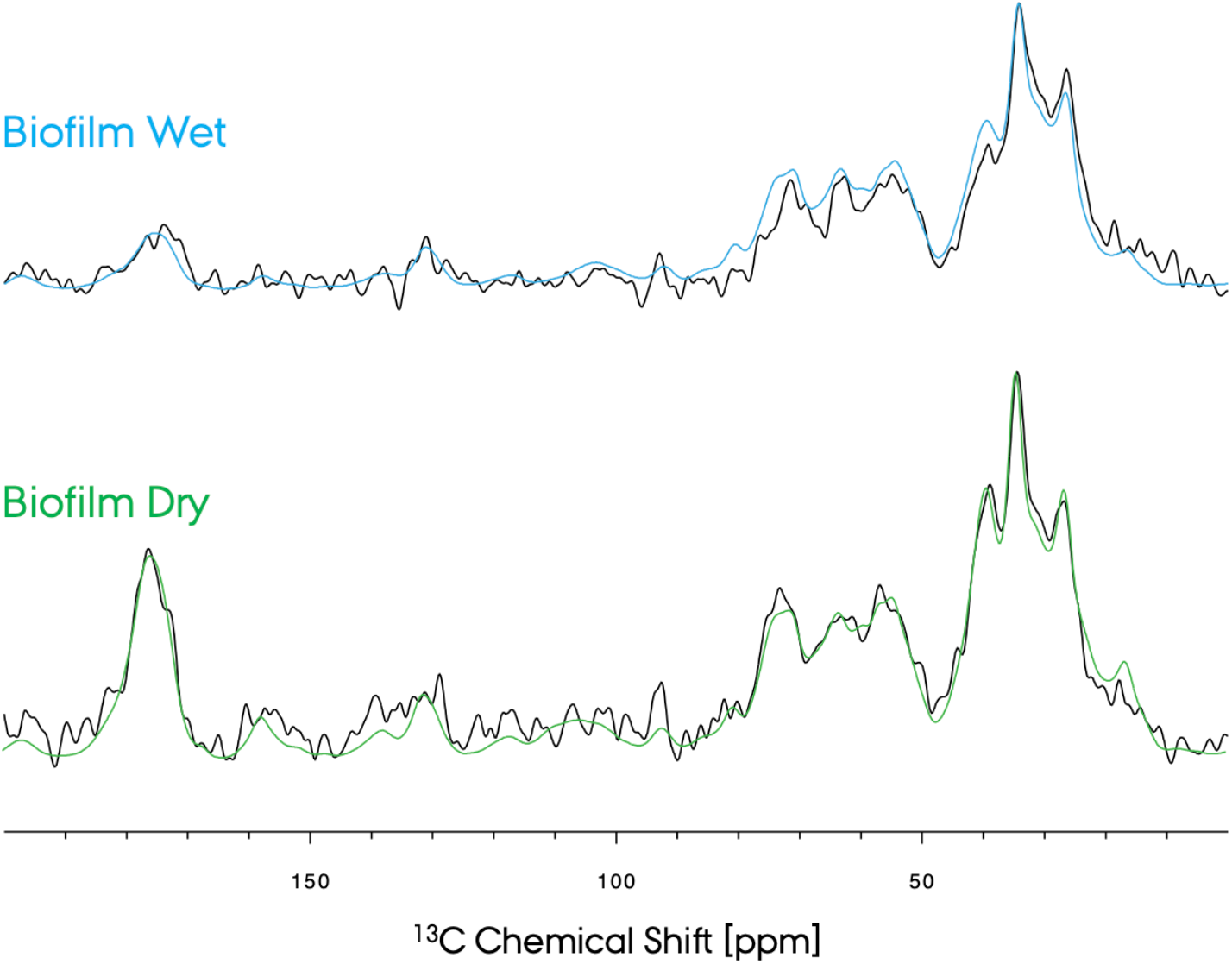
DNP enhancements obtained by comparing MW On/Off spectra for wet (blue) and dry (green) biofilm samples. The MW Off spectra are in black color for both samples. Overall, similar DNPO enhancements were observed for both samples.

**Figure SI 4:**
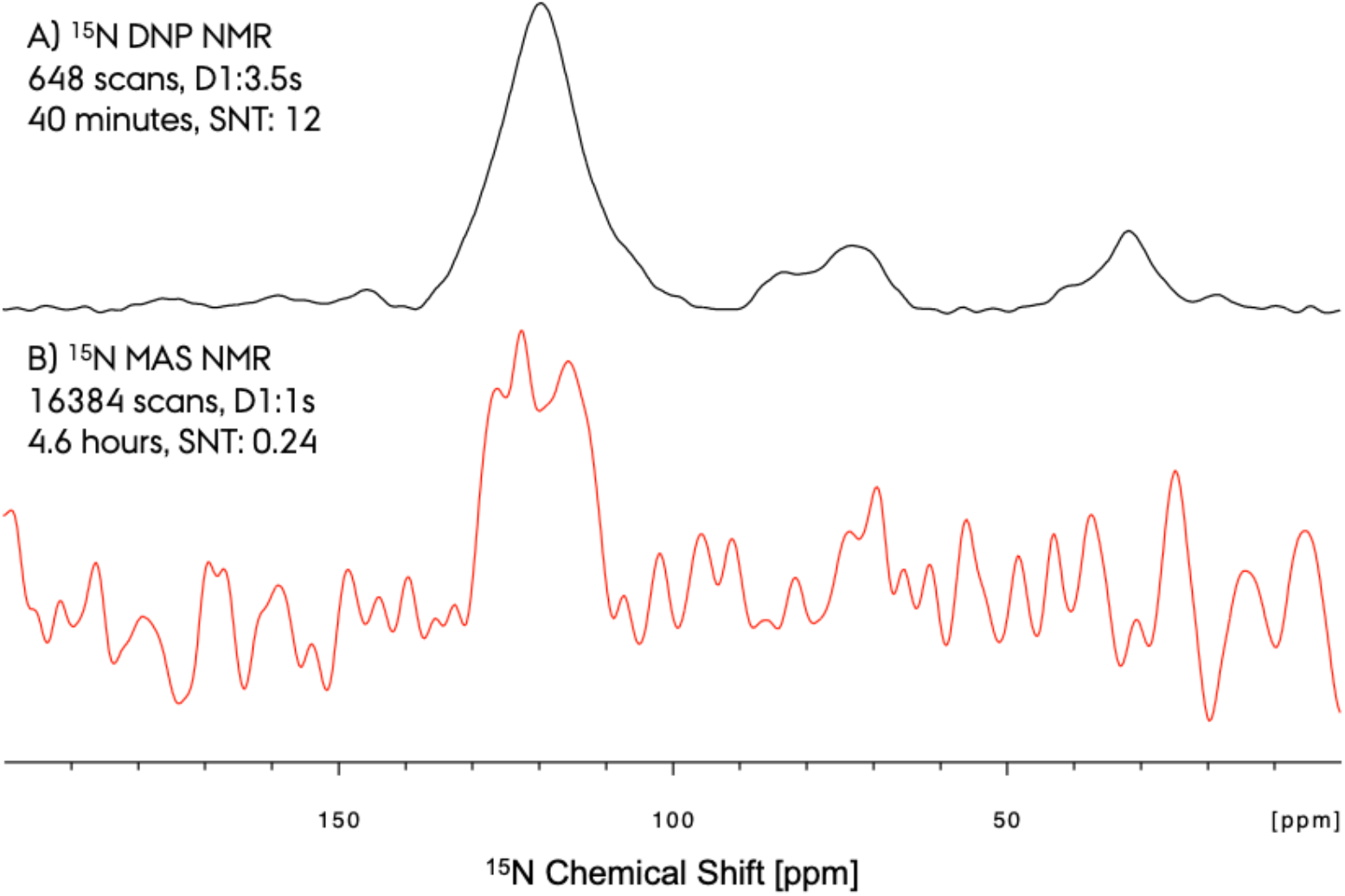
Experimental time reduction and sensitivity increase obtained for ^15^N via application of DNP ssNMR. Compared to the conventional room temperature spectra, the DNP ssNMR spectra is extremely more sensitive.

**Figure SI 5:**
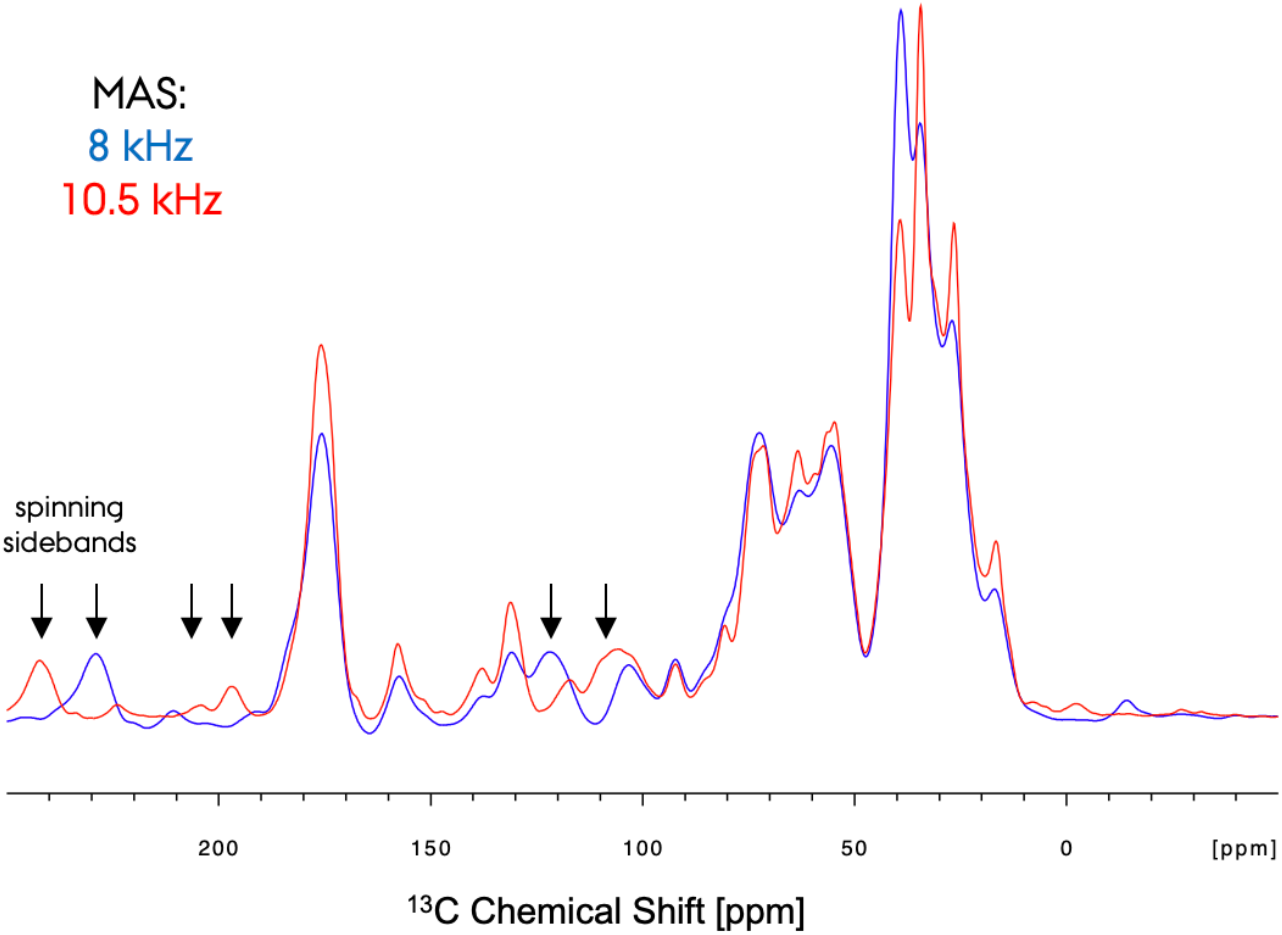
^13^C DNP ssNMR recorded at two different MAS frequencies of ∼8 (blue) and ∼10 (red) kHz. The spinning sidebands due to MAS are marked with arrows.

**Figure SI 6:**
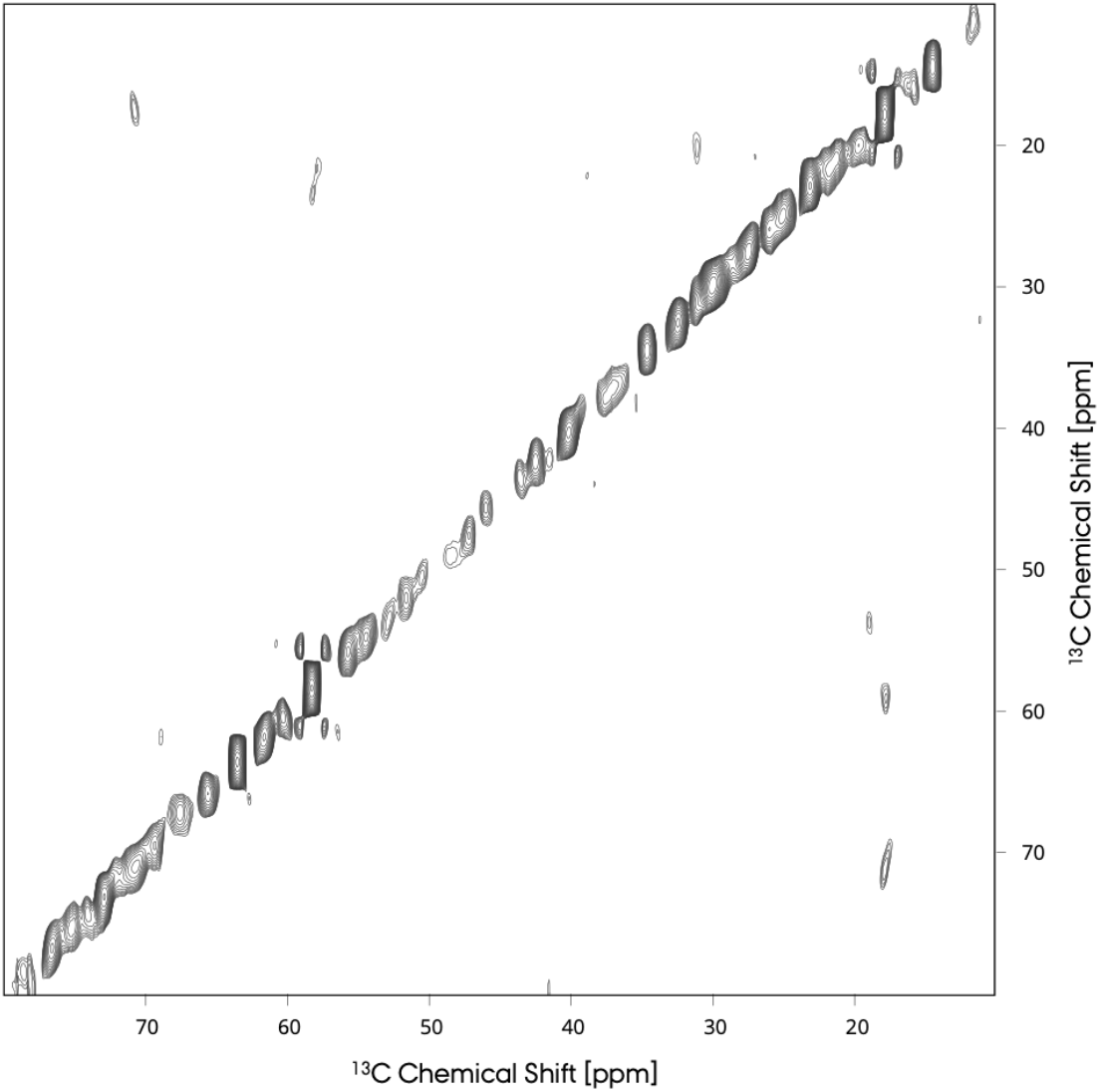
Conventional 2D ^13^C-^13^C PDSD recorded at room temperature in ∼200 hours with 20 ms mixing time, at 275 K and 10 kHz MAS. The 2D spectrum does not show any cross-peak due to the lack of sensitivity of the unlabeled biofilm sample.

